# Evaluation of the reliability and applicability of human unbound brain-to-plasma concentration ratios

**DOI:** 10.1101/2022.11.14.516429

**Authors:** Urban Fagerholm

## Abstract

**Background:** Blood-brain barrier permeability (BBB P_e_) and unbound brain-to-plasma concentration ratio (K_p,uu,brain_) are relevant parameters describing the brain uptake potential of compounds. BBB efflux by transporter proteins, mainly MDR-1 and BCRP, is an essential factor determining K_p,uu,brain_. K_p,uu,brain_-values are commonly estimated *in vivo* in rats and monkeys and predicted using *in silico* methodology. Such estimates can be used to predict corresponding human clinical values.

**Objective:** The objective of the study was to evaluate the reliability and applicability of human clinical K_p,uu,brain_-data for understanding and predictions of brain uptake in man.

**Methodology:** K_p,uu,brain_ in rats, monkeys and humans, measured and *in silico* predicted MDR-1 and BCRP substrate specificities and *in silico* predicted passive P_e_ were used for the analysis. *In silico* predictions were done using the ANDROMEDA by Prosilico ADME/PK-prediction software.

**Results and Discussion:** Rat and monkey K_p,uu,brain_-values were highly correlated (R^2=0.74; n=17). Based on this finding a correlation between rat and human K_p,uu,brain_ was expected. However, no correlation between rat and human K_p,uu,brain_ was found (R^2=0.01; n=13). There was no (as also anticipated) correlation between passive P_e_ and human K_p,uu,brain_ (R^2=0.04; n=16) and compounds with measured or predicted efflux did not have lower K_p,uu,brain_ than compounds without efflux. The compound with highest K_p,uu,brain_ in man (2.8) is effluxed and predicted to have high passive P_e_ and has no apparent efflux at the rat BBB. The MDR-1 substrate with highest K_p,uu,brain_ in rat (2.4) has very low K_p,uu,brain_ in man (0.15) is predicted to have high passive P_e_.

**Conclusion:** Results indicate that available human K_p,uu,brain_-data are too uncertain to be applicable for validation of predictions and understanding of clinical brain uptake of drugs and drug candidates.

## INTRODUCTION

Blood-brain barrier permeability (BBB P_e_) and unbound brain-to-plasma concentration ratio (K_p,uu, brain_) are relevant parameters describing the brain uptake potential of compounds (Summerfield et al. 2008; Syvänen et al. 2009; Chen et al. 2011; Sato et al. 2021; Loryan et al. 2022). BBB efflux by transporter proteins, mainly MDR-1 and BCRP, is an essential factor determining K_p,uu, brain_ (Chen et al. 2011; Sato et al. 2021; Loryan et al. 2022).

K_p,uu,brain_-values are commonly estimated *in vivo* in rats and monkeys and predicted using *in silico* methodology (Summerfield et al. 2008; Syvänen et al. 2009; Chen et al. 2011; Sato et al. 2021; Loryan et al. 2022). Provided that there are sufficiently strong correlations between preclinical estimates/models and human clinical values, human K_p,uu,brain_-values and brain uptake potential can be predicted using such estimates.

The objective of the study was to evaluate the reliability and applicability of human clinical K_p,uu,brain_-data for understanding and predictions of brain uptake in man.

## METHODOLOGY

K_p,uu,brain_ in rats, monkeys and humans, measured and *in silico* predicted MDR-1 and BCRP substrate specificities and *in silico* predicted passive P_e_ were used for the analysis. Sato et al. 2021 was the main source of rat, monkey and human K_p,uu,brain_-values and MDR-1 and BCRP substrate specificities. Additional rat and human K_p,uu,brain_-values were taken from Summerfield et al. 2008.

*In silico* predictions of passive P_e_ and substrate specificity for MDR-1 and BCRP (in cases where measured values were not available) were done using the ANDROMEDA by Prosilico ADME/PK-prediction software (Fagerholm et al. 2022).

## RESULTS & DISCUSSION

Rat and monkey K_p,uu,brain_-values were highly correlated (R^2=0.74; n=17; Figure 1). Based on this finding, a correlation between rat and human K_p,uu,brain_ was expected.

**Figure 1.**
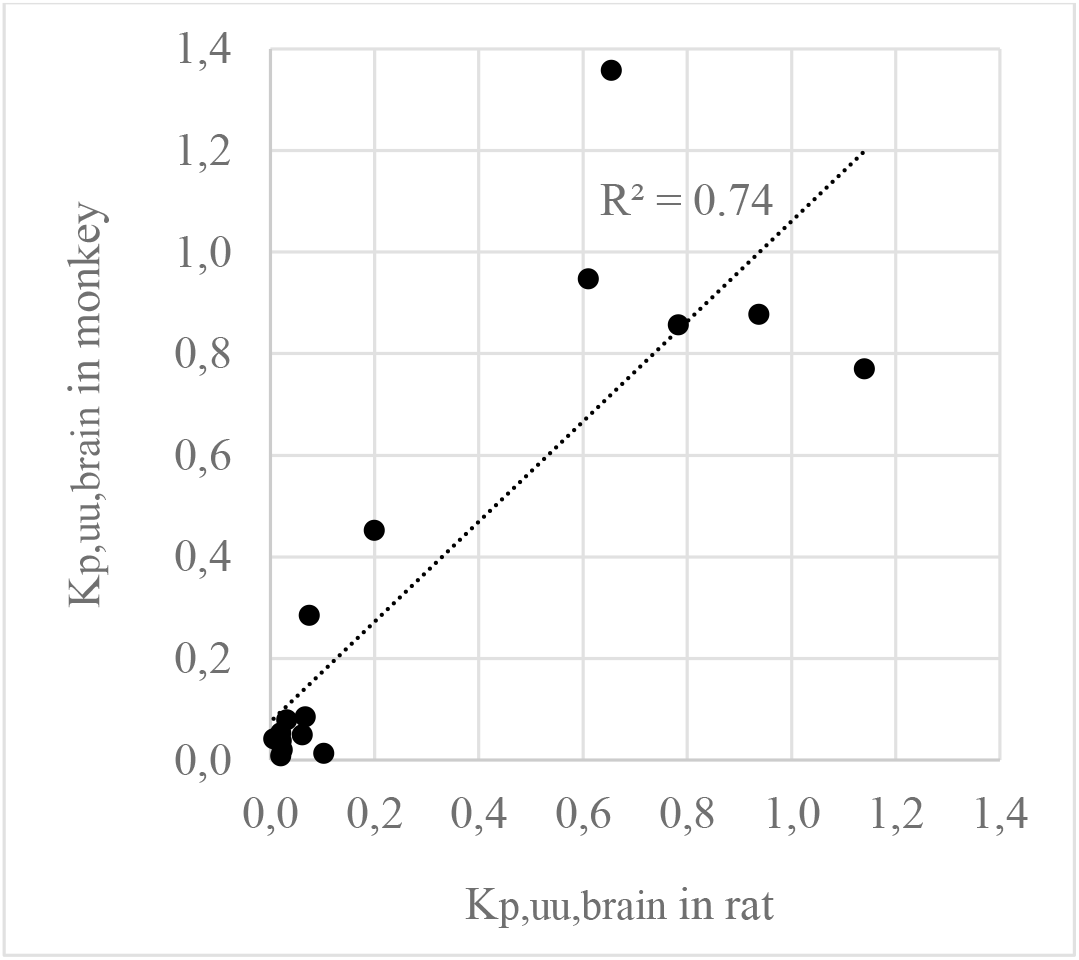
The correlation between rat and monkey K_p,uu,brain_-values (n=17; Sato et al. 2021).

In Sato et al. 2021 and Summerfield et al. 2008 there are rat and human K_p,uu,brain_-data for 21 compounds. There was, however, no correlation between rat and human K_p,uu,brain_ (R^2=0.01; n=13; Figure 2; using data from Sato et al. 2021, R^2=0.00 using data from Sato et al. 2021 and Summerfield et al. 2008).

**Figure 2.**
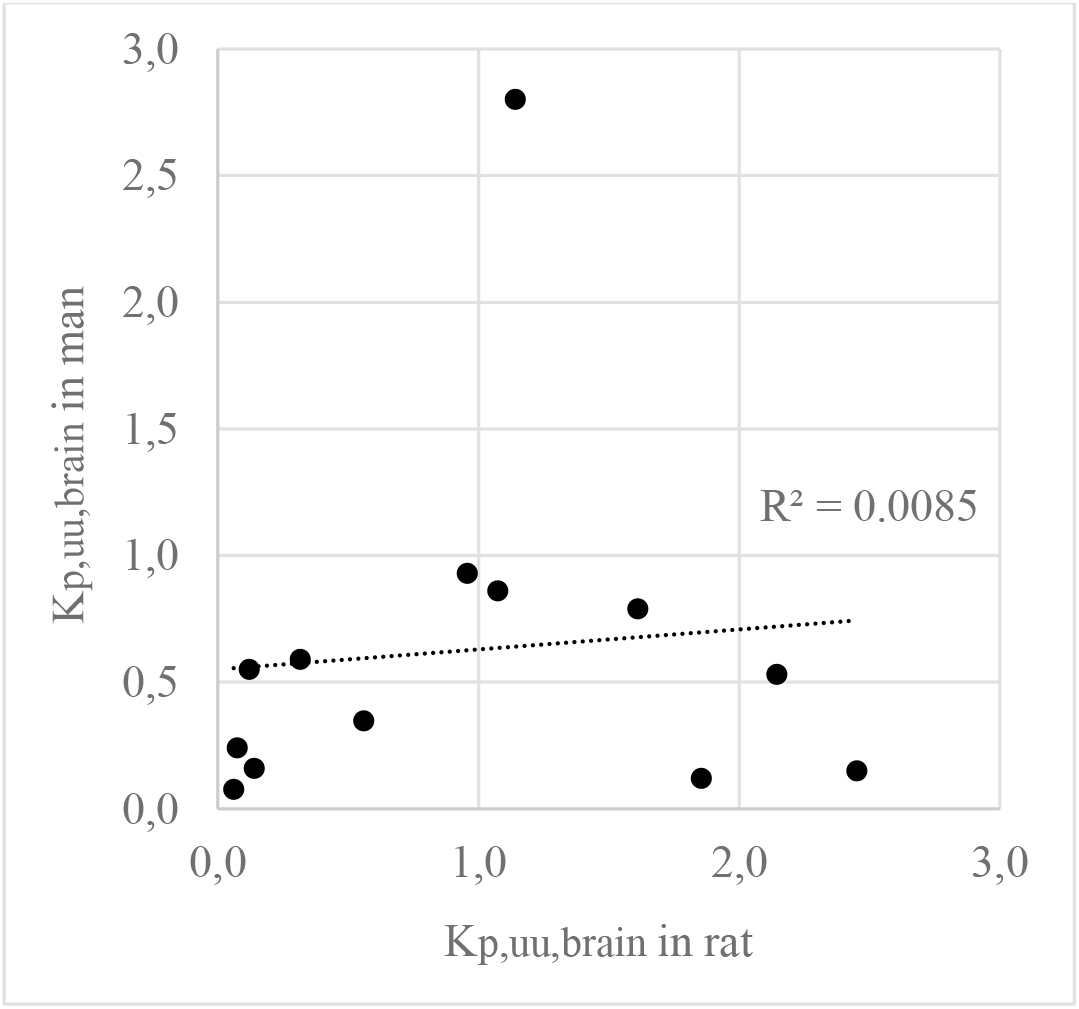
The correlation between rat and human K_p,uu,brain_ (n=13; Sato et al. 2021).

A higher dependency of efflux on K_p,uu,brain_ was anticipated at low P_e_, and therefore, a positive correlation between human passive P_e_ and K_p,uu,brain_ was expected. There was, however, no correlation between passive P_e_ and human K_p,uu,brain_ (R^2=0.04; n=16). Furthermore, compounds with measured or predicted efflux did not have lower K_p,uu,brain_ than compounds without efflux. The compound with highest K_p,uu,brain_ in man (lamotrigine; 2.8) is effluxed and predicted to have high passive P_e_ and has no apparent efflux at the rat BBB. The compound with the second highest K_p,uu,brain_ in man (flumazenil; 1.7) is effluxed and predicted to have high passive Pe. The MDR-1 substrate with highest K_p,uu,brain_ in rat (olanzapine; 2.4) has very low K_p,uu,brain_ in man (0.15) is predicted to have high passive P_e_. Cefotaxime is the compound with the lowest K_p,uu,brain_ in man (0.007) and it was predicted to have the lowest passive P_e_ (corresponding to less than 1 % oral uptake) and maybe BCRP-efflux (undetermined regarding BCRP-specificity).

At least 14 of the 16 compounds with human K_p,uu,brain_-estimates in Sato et al. 2021 had measured and/predicted MDR-1 and/or BCRP-efflux. 14 of them were also predicted to have high passive P_e_.

Results indicate that available human K_p,uu,brain_-data are too uncertain to be applicable for validation of predictions and understanding of clinical brain uptake of drugs and drug candidates. An explanation could be methodological differences. Human data were obtained post-mortem (reduced/no blood flow and transporter activity) and in PET-studies using radioligands (non-equilibrium; radioligand degradation), while animal data were commonly produced in microdialysis (equilibrium) studies. Uncertainties in measurements of unbound fractions in plasma/blood and brain tissue and laboratory variability (on average ca 2-fold differences between highest and lowest reported values in the rat) are also possible contributors.

The high correlation between rat and monkey K_p,uu,brain_ (R^2=0.74), and high degree of species homology for MDR-1; 85 % between rats and man and 93-97 % between monkeys and man; Syvänen et al. 2009) indicates that data obtained these two species are likely to correlate to corresponding true human values to a similar extent. A 15 % difference in MDR-1 homology and relatively high expression of MDR-1 and comparably low expression of BCRP in rats demonstrate that significant true K_p,uu,brain_-differences between these two species may also occur sometimes. Measurements and/or predictions of MDR-1 and BCRP-substrate specificities and ratios are useful for improving the certainty of human K_p,uu,brain_-predictions. The new 3-dimensional brainavailability-matrix (passive BBB P_e_-class *vs* brain binding-class + efflux/non-efflux and K_p,uu,brain_) is also applicable for a better overview (Fagerholm et al. 2022).

